# Nomadic ungulate movements under threat: Declining mobility of Mongolian gazelles in the Eastern Steppe

**DOI:** 10.1101/2023.02.05.526430

**Authors:** Philipp Mendgen, Nandintsetseg Dejid, Kirk Olson, Bayarbaatar Buuveibaatar, Justin M. Calabrese, Buyanaa Chimeddorj, Munkhnast Dalannast, William F. Fagan, Peter Leimgruber, Thomas Müller

**Affiliations:** Senckenberg Biodiversity and Climate Research Centre, Senckenberg Gesellschaft für Naturforschung, Senckenberganlage 25, 60325 Frankfurt, Germany; Department of Biological Sciences, Goethe University, Max-von-Laue-Straße 9, 60438 Frankfurt, Germany; Wildlife Conservation Society, Mongolia Program, Ulaanbaatar 14200, Mongolia; CASUS - Center for Advanced Systems Understanding (CASUS), Helmholtz-Zentrum Dresden-Rossendorf e.V. (HZDR), Untermarkt 20, 02826 Görlitz, Germany; Department of Ecological Modelling, Helmholtz Centre for Environmental Research-UFZ, Permoserstrasse 15, 04318 Leipzig, Germany; Department of Biology, University of Maryland, College Park, MD, USA; World Wide Fund for Nature, Mongolia Program Office, Ulaanbaatar 14200, Mongolia; Department of Biology, Mongolian National University of Education, 14191 Ulaanbaatar, Mongolia; Conservation Ecology Center, Smithsonian Conservation Biology Institute, National Zoological Park, 1500 Remount Road, Front Royal, VA 22630, USA

**Keywords:** Animal movement, GPS tracking, semipermeable barriers, traffic, ungulate

## Abstract

Increasing habitat fragmentation and disturbance threaten long distance movements of ungulates. While the effects of impermeable barriers on ungulate migrations have been well researched, quantitative evidence for gradual and long-term changes of mobility in response to anthropogenic disturbance remains relatively rare.

We investigated changes in movement behavior of Mongolian gazelle *Procapra gutturosa*, a nomadic ungulate species native to the Mongolian steppe. Using GPS tracking data collected from 62 gazelle individuals between 2007 and 2021, we quantified 16-day displacement distances for each individual as a metric for long-distance movements. We used generalized linear mixed models, generalized additive models and additive quantile mixed models to assess how anthropogenic and environmental factors affected gazelle movement behavior.

Long distance 16-day movements decreased significantly by up to 36 %, from 142 km in 2007 to 92 km in 2021. Changes in gazelle mobility were affected by the increasing number of vehicles in Mongolia, but could not be explained by concurrent changes in other environmental factors like temperature, precipitation or vegetation greenness that often drive ungulate migration behavior. Moreover, we found that gazelle movement decreased close to roads, and that gazelles stayed further away from roads during the snow-free season, when vehicular traffic likely is most intense.

Conserving landscape permeability is essential for maintaining populations of highly mobile species. Our study provides evidence for a gradual decline in gazelle mobility over fifteen years as a response to increasing anthropogenic impact. To date, the transportation infrastructure permeating the Eastern Steppe does not pose physical barriers, yet our findings suggest that increasing traffic volume may create semipermeable barriers to gazelle movement. As human activity is increasing throughout the Eastern Steppe, interactions between ungulates and vehicle traffic need to be closely monitored in order to identify, localize, and mitigate semipermeable barrier effects before landscape permeability is severely altered.

## Introduction

The movements of land mammals are diminishing worldwide due to anthropogenic disturbances that change animal behavior (Tucker et al., 2018). This effect is apparent across many taxa, influencing species directly and cascading through ecosystems as critical services provided by animal movements (e.g., seed dispersal) decline (Løvschal et al., 2017; Lundberg & Moberg, 2003). Among the world’s mammal species, ungulates are particularly affected due to their widespread reliance on long-distance-movement such as seasonal migrations, which enable them to access vital resources (Fynn & Bonyongo, 2011; Merkle et al., 2016) and to escape from unfavorable environmental conditions (Folstad et al., 1991; Monteith et al., 2011). Consequentially, far-ranging ungulate species, such as wildebeest (*Connochaetes taurinus*), bison (*Bison bison*), or saiga antelope (*Saiga tatarica*) have become increasingly threatened due to the development of anthropogenic barriers, habitat loss, and overhunting (Berger, 2004; Bolger et al., 2008). Mobility constraints caused by impermeable barriers are perhaps the most direct threat to long distance ungulate mobility (Harris et al., 2009; Sawyer et al., 2013). Where long distance movements are no longer possible, the abundance of migratory ungulate species often declines drastically, as has been observed in the past (Harris et al., 2009) and predicted for the future (Fryxell et al., 2004; Holdo et al., 2011). Even barriers that are not completely impermeable have the potential to change movement behaviors, thereby reducing the benefits of migration (Aikens et al., 2022; Sawyer et al., 2013). For such barriers, the degree of permeability may vary depending on the level of human activity, and mobility may become severely impeded once certain disturbance thresholds are crossed (Sawyer et al., 2020).

Longitudinal studies documenting the continuous and long-term decline of long-range movements are rare. Much of the existing research on reductions of ungulate mobility through anthropogenic barriers focused on changes in population abundance (Harris et al., 2009), spatial comparisons of mobility between affected and unaffected populations (Lendrum et al., 2012) or relied on modelling (Holdo et al., 2011). Some attempts at longitudinal studies have leveraged multi-year GPS data to investigate effects of impermeable barriers (Ito et al., 2013), avoidance of disturbed areas (Sawyer et al., 2020) or increased migration speed when moving though disturbed area (Sawyer et al., 2013).

We studied the effect of anthropogenic disturbance on the mobility of nomadic Mongolian gazelle *Procapra gutturosa* over the course of 15 years. Mongolian gazelles (hereafter gazelles) are far-ranging nomadic ungulates that primarily occur in the Gobi-Steppe ecosystem of Mongolia (Clark et al., 2006). They are iconic for their vast aggregations and long distance movements in what is considered the largest intact temperate steppe ecosystem worldwide (Batsaikhan et al., 2014). Previous research shows that individual gazelle are capable of roaming through up to 23,000 km^2^ per year, forming lifetime ranges of more than 100,000 km² (Nandintsetseg et al., 2019). This high mobility is crucial for the survival of gazelle, as areas of high-quality grasses and forbs occur dynamically in space and time (Mueller et al., 2008). As a result, the movements of gazelle appear to be highly irregular and unpredictable, with no strong fidelity to particular calving or winter ranges, no consistent migratory routes, and huge temporary aggregations of individuals in times of resource scarcity (Olson et al., 2009, 2010).

Mongolia is currently experiencing dramatic socioeconomic transformations and rapid economic growth as it is exploiting previously untapped, rich mineral and fossil fuel deposits (Batsaikhan et al., 2014; Reading et al., 2006). These societal shifts are accompanied by the expansion of linear infrastructure networks throughout the steppe (Batsaikhan et al., 2014). Our study focuses on gazelles moving in Eastern Steppe within the Dornod and Sukhbaatar provinces, which represent an important stronghold of the species’ population. Here, road infrastructure is comprised of unfenced paved and dirt roads that do not pose much of a physical barrier to ungulate movement, but create extended corridors of degraded vegetation and human disturbance (Keshkamat et al., 2013). However, motorized traffic in the region has been steadily increasing (National Center for Road Transport, 2022). In such an environment, road infrastructure may be semi-permeable to ungulate movement, and barrier effects may increase with increasing human activity (Jacobson et al., 2016; Sawyer et al., 2013). Our aim was to investigate whether the mobility of gazelle in the Eastern Steppe had changed over the last 1.5 decades, and if so, whether such change could be attributed to anthropogenic disturbance.

We analyzed changes in gazelle displacement distances over 15 years. We assessed displacement over long timescales (average 16-day displacement and the 90% quantile of 16-day displacement) as human disturbance may alter movement tortuosity rather than speed (Tucker et al., 2018), and encounter effects could be cumulative (Eftestøl et al., 2021). To identify potential drivers of longitudinal changes in gazelle movement, we assessed whether changes in vehicle numbers in Mongolia (a proxy for traffic) affected gazelle mobility, while accounting for the effect of environmental drivers such as temperature, precipitation, snow cover, and vegetation greenness, which have been shown to greatly influence ungulate migration (Holdo et al., 2009; Laforge et al., 2021; Mason et al., 2014; Merkle et al., 2016). We could not directly test for effects of traffic volume and road development since longitudinal data on traffic and road construction in the Eastern Steppe were not available. We hypothesized that increases in traffic and road construction change landscape permeability in the Eastern Steppe, thereby reducing long-distance mobility of gazelles. Because traffic is generally lower in winter and gazelle may be more tolerant of anthropogenic structures during this season (Ito et al., 2013), we also hypothesized that long-term mobility declines are more apparent in summer. To corroborate effects of road infrastructure on gazelle mobility, we tested whether gazelle displacement distances decreased near roads using static spatial data of those features. We also tested whether gazelle approached roads closer in winter than in summer, and whether this might be more apparent in gazelle that stayed generally closer to human disturbance compared to gazelle that stayed farther away overall.

## Methods

### Movement data of gazelle

We compiled existing GPS movement data originating from capture and collaring efforts between 2007 and 2021. All captures followed the standard protocols approved by the Ministry of Environment and Green Development in Mongolia (see details in D. Nandintsetseg et al., 2019; Olson et al., 2014). This data set included tracking data of 62 gazelle individuals with varying fix rates (see appendix A1 for details). Across these fix rates, the best feasible interval for the joint analysis of gazelle displacements (the straight-line distances between observations) over long timescales was 16-day intervals. We performed all analyses using R (R Core Team, 2021).

### Environmental and human disturbance data

We obtained environmental covariates from the Land Processes Distributed Active Archive Center (LPDAAC), extracting monthly temperature, snow cover, and normalized difference vegetation index (NDVI) data for the years 2007 - 2021, obtained at 0.05-degree resolution by Terra MODIS (Didan, 2015; Hall & Riggs, 2015; Wan et al., 2015). From the Climate Hazards Group InfraRed Precipitation with Station data (CHIRPS) data archive (Funk et al., 2014), we acquired monthly precipitation data for the same time periods, also at 0.05-degree resolution.

We acquired yearly data on the number of registered vehicles in Mongolia between 2012 and 2021 (National Center for Road Transport, 2022). Data on registered vehicles in 2007-2011 were not available, however we obtained data on the number of imported vehicles during that time and used them to calculate the number of vehicles in Mongolia in that period (assuming that imported vehicles were registered in the year of import, and that no vehicles were produced in Mongolia). To measure the proximity of gazelles to road infrastructure we accessed road maps from the Administration of Land Affairs, Geodesy, and Cartography of Mongolia (ALAGaC, Agency of Land Administration and Management Geodesy and Cartography Mongolia, 2022) and from OpenStreetMap (OpenStreetMap Contributors, 2022). These spatial data provide a snapshot of the current distribution of disturbances throughout the steppe.

For each displacement step, we calculated mean temperature, precipitation, snow cover and NDVI, as well as the mean distance to the nearest road. We also recorded the year, month, and coordinates of the first position in each step, as well as the unique identifier for each individual gazelle. We classified each step into a snow-free season (April – October) or winter season (November – March). To assess long distance displacements, we determined whether any step belonged to the 10% of longest steps moved during each season per year, over all gazelles.

### Assessing longitudinal effects on displacement

We assessed the effect of year on gazelle displacement by building linear mixed models for the winter and snow-free seasons, using displacement distance as the response variable and year as the only predictor. To determine whether long distance movement was especially affected, we repeated the analysis using only the longest 10% of displacement steps. To assess potential anthropogenic drivers, i.e., the effect of increasing human activity while correcting for potential environmental changes, we built a second set of linear mixed modes using the number of vehicles (in hundred thousand), temperature (in °C), NDVI, precipitation (in mm), and snow cover (in %) as predictors. Models for the snow-free season did not incorporate snow cover. All models included a random intercept for gazelle individual to account for interindividual differences.

We computed Cullen and Frey graphs via the `fitdistrplus` package (Delignette-Muller & Dutang, 2015) to identify suitable transformations of the response variables, after which we cube-root transformed displacement in all models. We tested whether including a quadratic term of NDVI improved the predictive power of the model as measured by Akaike’s Information criterion (AIC; Akaike, 1974), as gazelles might prefer intermediate values of NDVI (Mueller et al., 2008). Subsequently we added a quadratic term for NDVI in all models that incorporated NDVI. We tested for correlation between predictor variables by calculating pairwise Pearson correlation coefficients. Temperature and snow cover were always highly correlated (ρ > |0.6|), so we included one of these parameters at a time and calculated the AIC to determine the parameter with the highest predictive power, and consequently kept temperature in the model while discarding snow cover (see appendix A2).

We assessed model assumptions using the `DHARMa` package (Hartig, 2021) (see appendix A3). In the presence of influential outliers or heteroscedasticity, we refitted the respective models as robust linear mixed effect models using the rlmer function in the `robustlmm` package (Koller, 2016). Because rlmer does not compute p-values, we used the t-statistics of the “robust” rlmer model together with the Satterthwaite degrees of freedom of the “normal” lmer model to determine significance (see e.g. Geniole et al., 2019; Gómez et al., 2022). We tested for the presence of spatial autocorrelation (SAC) in the model residuals using the test for Moran’s I in the `ape` package (Paradis & Schliep, 2019). In case we detected significant SAC, we employed Moran’s eigenvector maps following Bauman et al. (2018) to reduce SAC (see appendix A4). We calculated the partial R^2^ attributed to each model covariate using the `partR2` package (Stoffel et al., 2021) and computed back-transformed predicted marginal means with 0.95 confidence intervals for each predictor using the `ggeffects` package (Lüdecke, 2018) and the `emmeans` package (Lenth, 2021). Finally, we refitted all models without using variable transformations, robust models or autocorrelation corrections to ensure that significant trends did not arise as artifacts of those additions (see appendix A5).

### Assessing the effect of anthropogenic disturbance on displacement

We modelled the effect of proximity to roads on average 16-day gazelle displacement via generalized additive models, using the `mgcv` package (Wood, 2011). We employed a Gaussian Location-Scale Model via the “gaulss” family to achieve unbiased p-values in the presence of heteroscedasticity. As in the previous analyses, we cube-root transformed the response to achieve normally distributed residuals. Predictors included thin plate regression splines of the mean and standard deviation of the distance to the nearest road as well as random intercepts for gazelle individuals and years. We assessed spatial dependence in the model residuals as described previously and accounted for it by including a smoothed interaction term of the spatial coordinates at the start of each displacement step. We also tested whether an interaction between distance to road and season improved model fit using AIC (see appendix A6). Additionally, we log10-transformed the distance to roads as we were most interested to model potential effects close to the source of disturbance. Because roads might often be located close to other sources of human disturbance, we excluded displacement steps during which gazelles were observed within 10 km of a settlement, railway or oil extraction site from this analysis (353 of 1903 data points), using railway maps digitized from google earth imagery (Google Earth, 2022) and spatial data on the location of settlements and oil wells from ALAGaC (Agency of Land Administration and Management Geodesy and Cartography, 2022) and the environmental department of Dornod province (Department for Environment of Dornod Aimag, 2015).

### Assessing seasonal effects of disturbance

To test the effect of season on the proximity of gazelles to roads, we calculated the average distance of each gazelle to the nearest road during both the snow-free and winter season of each year. We tested the effect of season and year on the 0.1, 0.25, 0.5, 0.75 and 0.90 quantile of distances to roads in an additive quantile mixed model framework using the `aqmm` package (Geraci, 2019), including a random intercept to account for differences between individuals. Additive quantile mixed models produce robust results in the presence of non-normally distributed or heteroscedastic errors (Geraci, 2019; Yirga et al., 2021).

## Results

### Longitudinal effects on gazelle displacement

Over the 15-year study period, we detected a significant decrease in average gazelle movements at 16-day intervals during the snow-free season (β = -0.028, t = -3.025, p = 0.004), but not during winter (β = -0.001, t = -0.132, p = 0.895). During the snow-free season, the estimated marginal mean of 16-day displacement decreased from 44.4 km [0.95 ci: 38.6; 51.1] in 2007 to 31.3 km [27.3; 35.3] in 2021, constituting a decline by 29.5 % (13.1 km) over 15 years (Figure 2a). When including only the longest 10% of gazelle movements per year, we detected significant longitudinal decreases during both the snow-free (β = -0.050, t = -5.602, p < 0.001) and winter seasons (β = -0.034, t = -2.889, p = 0.006). In the snow-free period, the estimated marginal means of 16-day long distance displacement decreased from 142.2 km [128.8; 155.7] in 2007 to 91.7 km [84.6; 99.9] in 2021, constituting a decrease by 35.5 % (50.5 km) over 15 years (Figure 2c). In winter, the estimated marginal means of 16-day long distance displacement decreased from 119.1 km [103.8; 135.8] in 2007 to 87.5 km [77.9; 98.0] in 2021, decreasing by 26.5 % (31.6 km) (Figure 2d).

**Figure 1:**
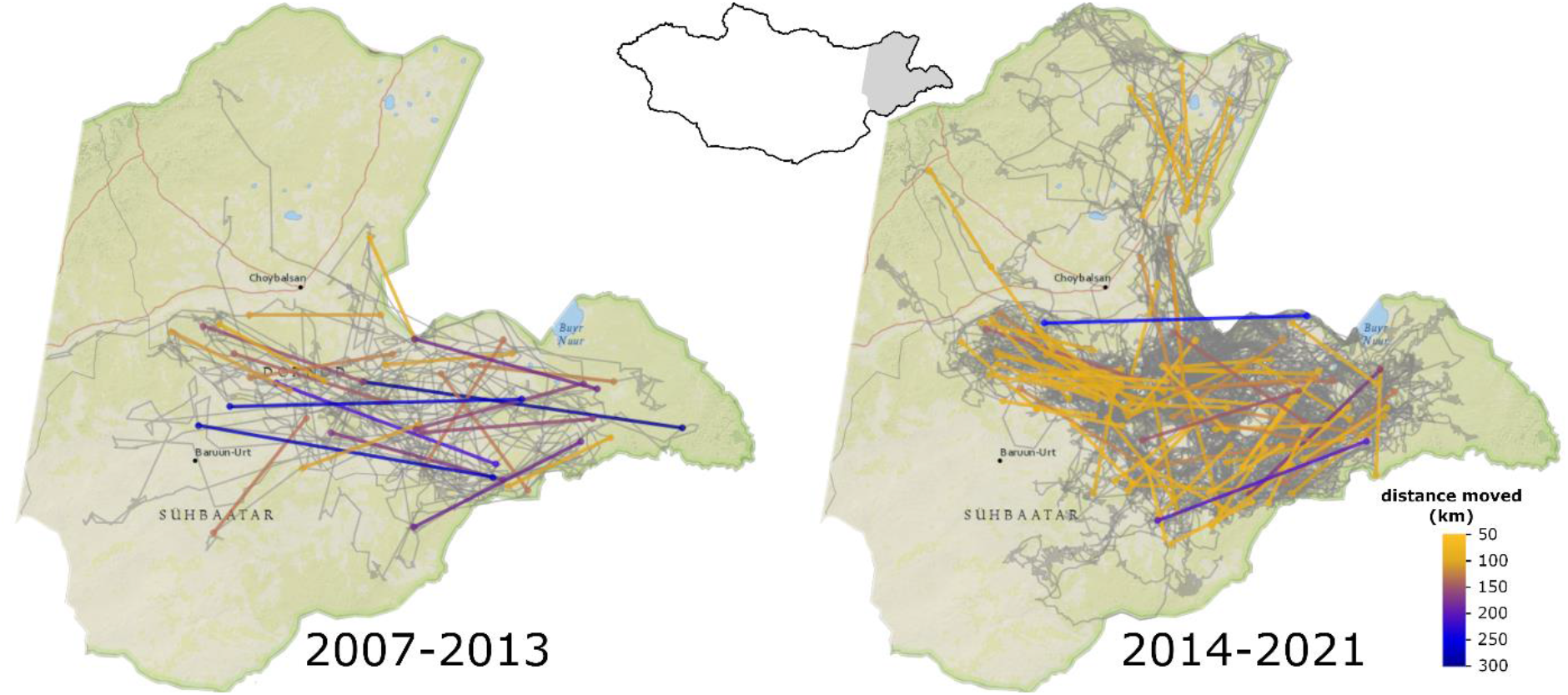
Gazelle long-distance displacements in summer as straight line 16-day distances, on top of actual gazelle movement paths (in grey). The background depicts the Dornod and Sukhbaatar provinces of Mongolia. This map was created via the R package leaflet, background tiles © Esri – National Geographic, Esri, DeLorme, NAVTEQ, UNEP-WCMC, USGS, NASA, ESA, METI, NRCAN, GEBCO, NOAA, iPC.

**Figure 2:**
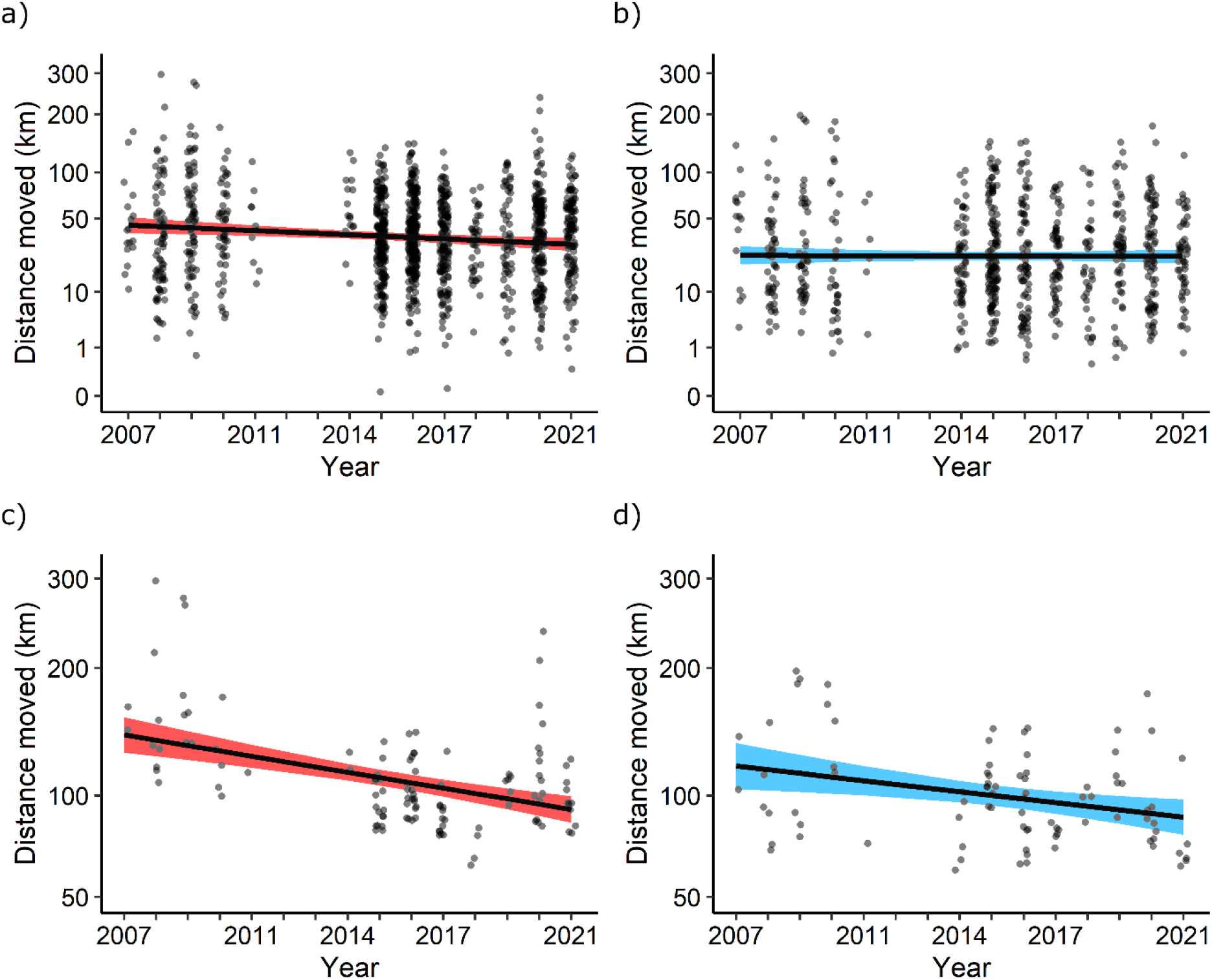
Effect of year on average and long-distance gazelle displacements. The predicted marginal means of the effect of year on the back-transformed average (a, b) and long-distance (c, d) 16-day displacements during the snow-free (a, c) and winter seasons (b, d) are depicted with 95% confidence bands. Average displacements decreased strongly over time during the snow-free season, while long-distance displacements decreased strongly during both seasons.

### Potential drivers of declines in mobility

The increasing number of vehicles in Mongolia negatively affected average gazelle mobility in the snow-free season (β = -0.050, t = -3.254, p = 0.002), and long-distance mobility of gazelles in both the snow-free (β = -0.078, t = -4.461, p < 0.001) and winter season (β = -0.049, t = -2.358, p = 0.024), while accounting for concurrent changes in environmental factors and inter-individual differences. Only average winter mobility was not significantly affected by number of vehicles (Figure 3). During the study period, the number of vehicles in Mongolia increased by 373.6 %, from 301,195 vehicles in 2007 to 1,125,385 vehicles in 2021. Concurrently, the estimated marginal means of average gazelle mobility decreased from 45.9 km [39.7; 52.7] to 31.9 km [27.5; 36.3] during the snow-free season, constituting a decrease of 14.0 km (30.6 %). For long-distance mobility, the estimated marginal means decreased from 136.6 km [120.6; 154.0] to 91.1 km [82.9; 100.5] during the snow-free season, constituting a decrease of 45.5 km (33.3 %); and from 113.4 km [98.6; 129.6] to 86.9 km [76.8; 98.0] during winter, constituting a decrease of 26.4 km (23.3 %). Temperature, NDVI, and precipitation did not significantly affect gazelle displacement in any model (Figure 3). The snow-free model explained 4.6 % of the variation in average gazelle displacement, with 1.2 % attributed to the vehicle effect. For long-distance mobility, the snow-free model explained 25.8 % of the variation in average gazelle displacement, with 16.8 % attributed to the vehicle effect; whereas the winter model explained 13.7 %, with 7.4 % attributed to the vehicle effect. See appendix A5 for model summary tables.

**Figure 3:**
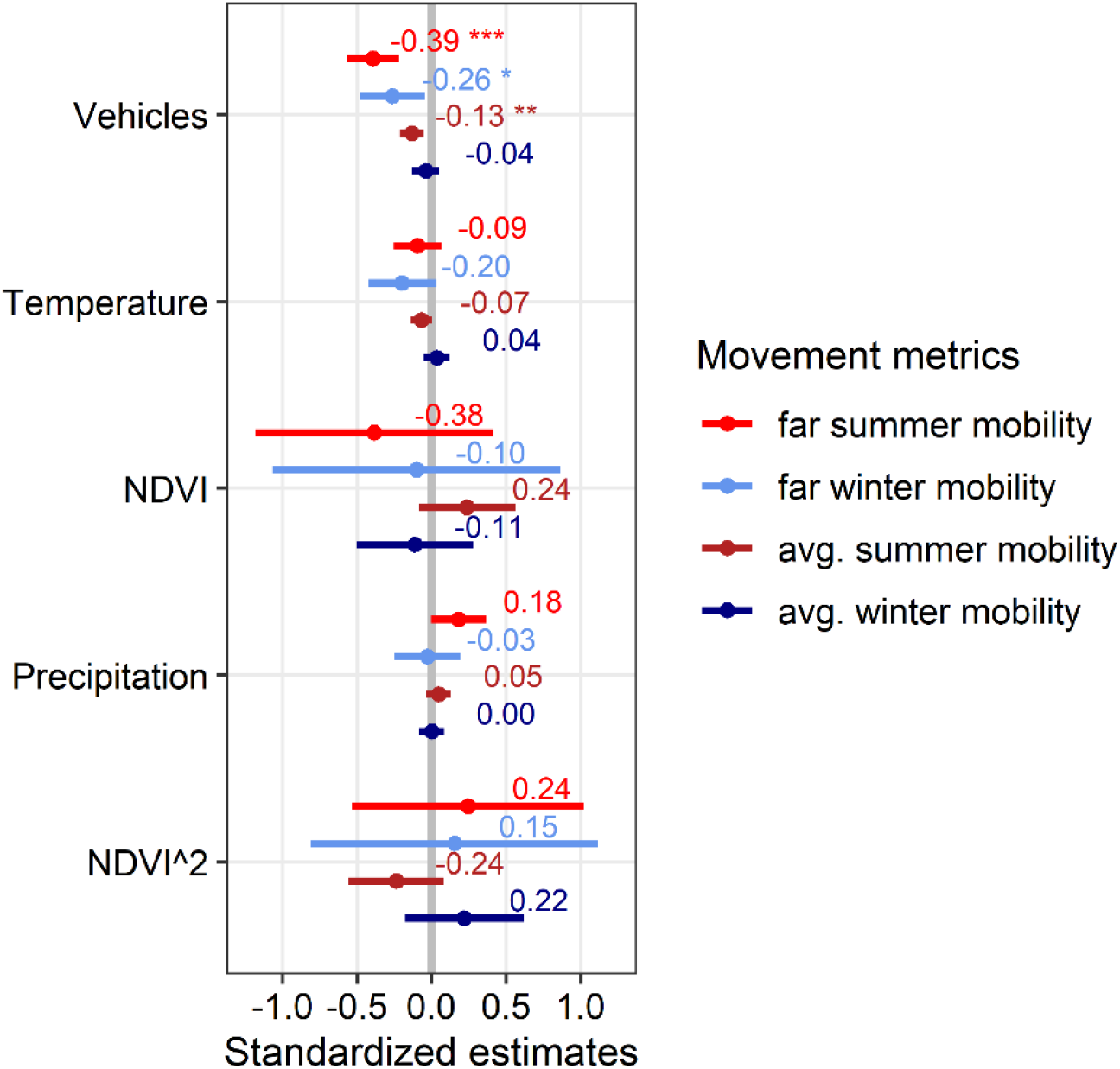
Standardized estimates of potential drivers of gazelle mobility for all constructed models. Vehicle number significantly affected gazelle mobility in all models except for average winter mobility, while no other predictor affected mobility. Standardized effect sizes were higher during the snow free season and for long-distance movement, compared to the winter season and average movement.

### Effects of road infrastructure on movement

The average distance to the nearest road significantly affected gazelle displacement (edf = 3.676, Chi.sq = 73.291, p < 0.001), while controlling for differences between individuals and years. Gazelle movements decreased in close proximity to roads, averaging 43.6 km [32.7; 58.1] at 10 km from the nearest road, but only 27.1 km [20.9; 35.1] while on average 1 km from it (Figure 4). The model explained 15.8 % of the residual deviance (10.6 % when excluding distance to road). There was no support for an interaction between distance to road and season. See appendix A5 for model summary tables.

**Figure 4:**
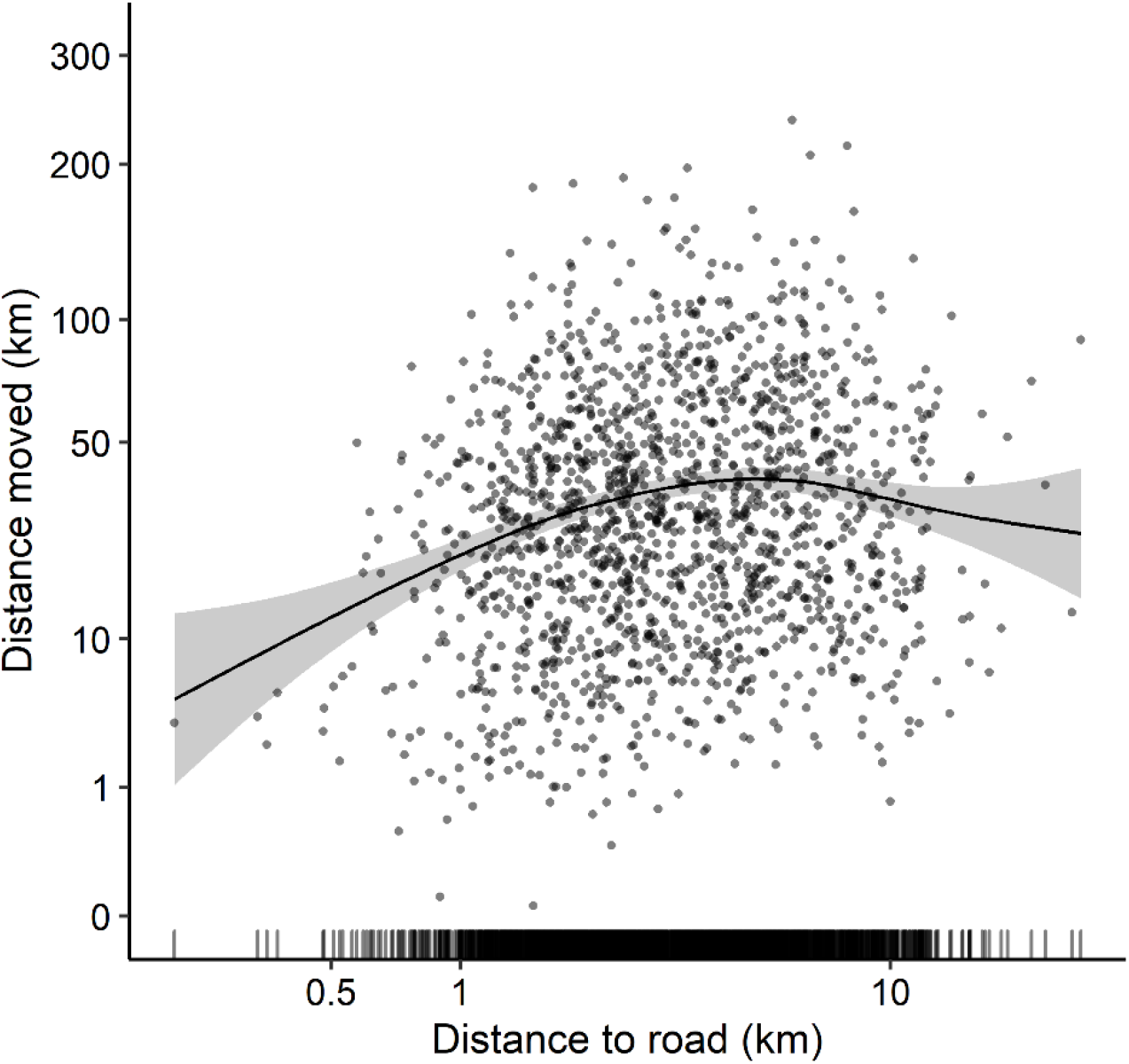
GAM response curve of the effect of distance to roads on 16-day displacement distances. Axis values are back-transformed to kilometers. Displacement distances decreased close to roads but stayed relatively constant once gazelles were several kilometers away from it.

### Seasonal differences of disturbance effects

Season significantly affected the 0.1, 0.25, 0.5 and 0.75 quantile of gazelle distances to roads. In winter, the estimated median distance to roads was 512.2 m (SE = 207.9, t = -2.463, p = 0.016) closer than the snow-free estimate of 3352 m, constituting a decrease of 15.3 % (Figure 5). The estimated 0.1 quantile of distances decreased by 261.4 m (SE = 101.2, t value = -2.584, p = 0.011) compared to the snow-free estimate of 1805.4 m (14.5 %), while the estimated 0.25 quantile of distances decreased by 438.6 m (SE = 148.2, t value = -2.960, p = 0.004) compared to the snow-free estimate of 2436.4 m (18 %). The 0.75 quantile of distances decreased by 770.7 m (SE = 210, t value = -3.670, p < 0.001) compared to the snow-free estimate of 4683 m (16.4 %). Year did not affect distances to roads in any quantile. See appendix A5 for model summary tables.

**Figure 5:**
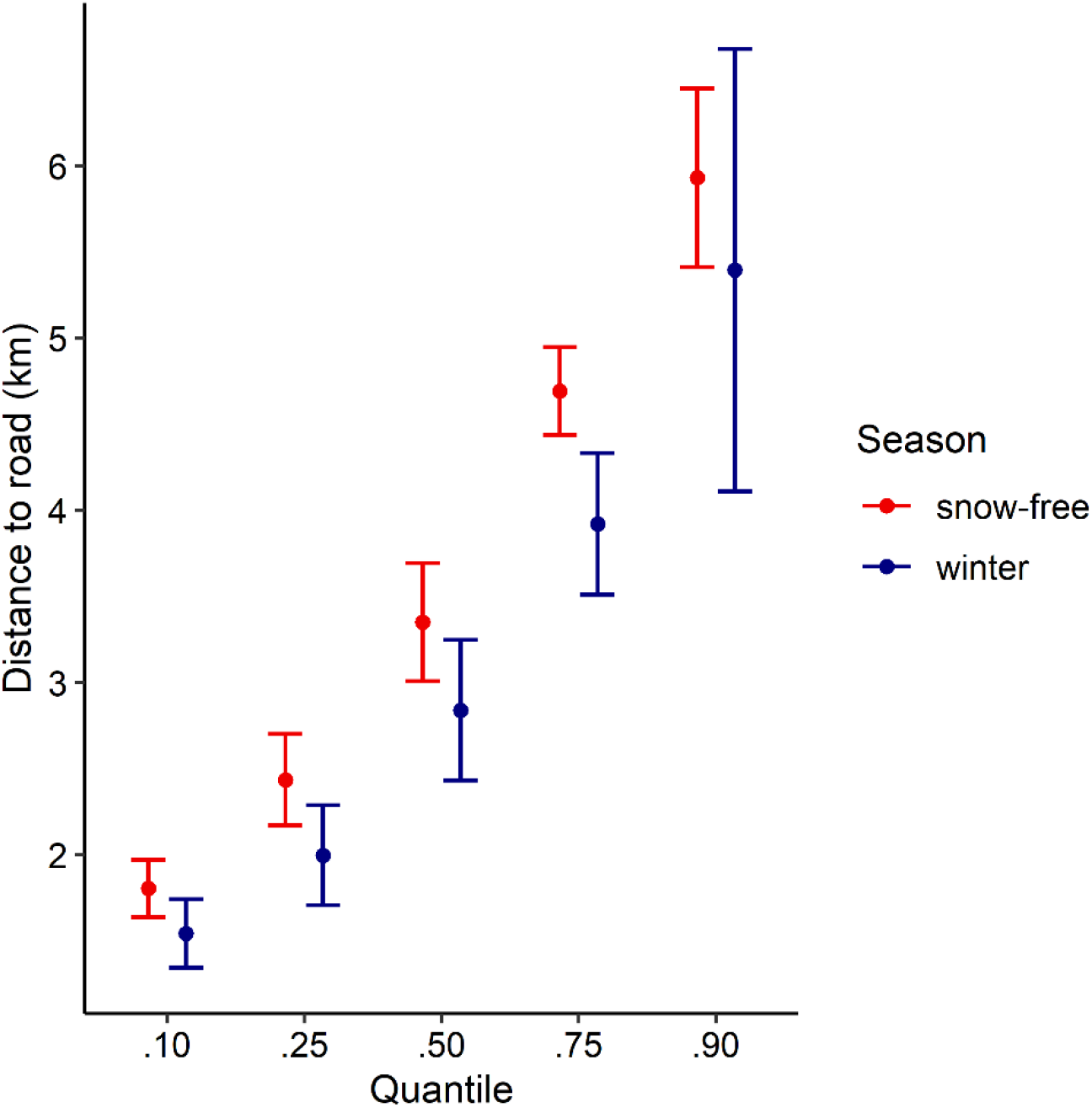
Seasonal differences in the distance of gazelles to the nearest road for multiple quantiles, with 95% confidence intervals. Most gazelles moved significantly closer to roads in winter, only gazelles that stayed very far from such disturbances did not.

## Discussion

We identified a decline of up to 13.1 km (29.5%) in the average and up to 50.5 km (35.5%) in the long-range movement behavior of gazelle covering a period of 15 years in a context of increasing anthropogenic disturbance. Decreases in gazelle mobility were correlated with the increasing number of vehicles and the presence of roads. Declines were more pronounced during the snow-free period, and gazelles displayed an increased avoidance of roads during that time. Our findings suggest that gazelle mobility in the Eastern Steppe decreased in response to human disturbance, potentially due to increases in road development and traffic.

Our study may be among the first to quantify gradual declines in ungulate movement distances over time. Decreases in ungulate migration have frequently been observed following the construction of anthropogenic barriers, which have affected migration propensity and migration routes in many ungulate populations, often culminating in the complete cessation of migratory behavior (Harris et al., 2009; Xu et al., 2021). Past research has collated quantitative evidence for the loss of ranges (Holdo et al., 2011; Said et al., 2016), routes (Berger, 2004) and abundances of migratory ungulate populations in response to anthropogenic development (Bolger et al., 2008; Holdo et al., 2011; Said et al., 2016). Previous research has also compared current and historic records of ungulate migration distances (Harris et al. 2009) and found that mammal movements decreased in areas of high human footprint using a space-for-time approach (Tucker et al. 2018). However, long-term individual-based data sets remain scarce but are urgently needed to determine how anthropogenic effects change ungulate behavior over time (Kauffman et al., 2021; Xu et al., 2021). Gradual effects of anthropogenic barriers have been observed in migratory mule deer (*Odocoileus hemionus*) where the increasing intensity and size of energy development caused detours in migration routes and affected migration timing (Aikens et al., 2022; Sawyer et al., 2013, 2017).

### Traffic and transportation infrastructure may affect gazelle mobility

Gazelles in our study may have responded to increases in traffic volume creating a disturbance barrier, rather than to a physical barrier created by roads. Contrary to many European or North American study systems, roads in the Eastern Steppe usually are dirt roads that likely do not pose impermeable physical barriers (Keshkamat et al., 2013; Yang et al., 2018). In addition, vehicle traffic has been observed to induce road avoidance behaviors in a wide range of species through disturbance cues such as noise or movement (Fahrig & Rytwinski, 2009; Jaeger et al., 2005), creating semi-permeable barriers whose effect could be mediated by the level of disturbance (Alexander et al., 2005; Sawyer et al., 2013). Such effects have also been observed near unpaved roads with low traffic volumes (D’Amico et al., 2016). Ungulate species known to avoid roads specifically due to vehicle traffic include red deer (*Cervus elaphus*) and wild boar (*Sus scrofa*) (Gagnon et al., 2007; Thurfjell et al., 2015), while others like pronghorn and Tibetan antelope (*Pantholops hodgsonii*) displayed increased vigilance behavior at high levels of traffic, suggesting they perceived cars as a threat (Gavin & Komers, 2007; Lian et al., 2011). Vehicle avoidance in gazelle may be exacerbated by the frequent use of cars and motorcycles for hunting. The plain landscape of the steppe makes wildlife easily accessible by vehicle, and both legal and illegal hunting frequently occur (Olson et al., 2014). During soviet times, gazelles were commercially hunted in large numbers, sometimes using trucks and automatic weapons (Lhagvasuren & Milner-Gulland, 1997). We observed gazelles to be shy to cars, initiating flight responses if a single car passed by them at several hundred meters distance and low speed (pers. comm. N. Dejid), although in some other regions to the south-west outside our study area gazelles may already show signs of habituation to cars where traffic volumes are high but hunting is less frequent (pers. comm. K. Olson). Hunting also affected road interactions in moose, increasing road avoidance during hunting season and in individuals that experience high hunting pressure (Paton et al., 2017). Road effects are known to be more severe for far-moving species because they encounter linear features frequently (Fahrig & Rytwinski, 2009; Jakes et al., 2018), which may also explain why our proxy for traffic explained much more variation in long-distance movement than in average movement.

### Seasonal differences in gazelle reactions to anthropogenic disturbance

We observed that declines in average and long-distance mobility were less strong in winter than during the rest of the year, while gazelles also stayed closer to roads in winter. This reduced avoidance of anthropogenic infrastructure during winter is consistent with findings of Kaczensky et al. (2006) and Ito et al. (2013), who reported that both gazelle and Asiatic Wild Ass (*Equus hemionus*) moved closer to anthropogenic barriers during that time. If the observed reductions in gazelle mobility are caused by traffic, the decreased human activity during the harsh Mongolian winter could explain why reductions of gazelle mobility were less pronounced during that time. Alternatively, the need to escape inhospitable environmental conditions such as snowstorms, deep snow cover, or lack of forage could have temporarily outweighed the avoidance of anthropogenic features in gazelle during winter. In other ungulates, pronghorn and mule deer avoided anthropogenic disturbance in winter, but tolerance for anthropogenic disturbance increased during migration; i.e. when there was a need to move through disturbed area (Sandoval Lambert et al., 2022).

### Consequences of decreasing mobility for nomadic ungulates

Decreased locomotion capacity can have drastic consequences for population levels of ungulates (Harris et al., 2009), causing events of high mortality in many migratory populations due to lacking access to vital resources (Williamson et al., 1988). Nomadic ungulate populations, such as gazelle, may be more resilient to anthropogenic impacts than migratory populations due to their high plasticity in route and sites use, yet they are also difficult to protect due to their far-ranging movements (Nandintsetseg et al., 2019), and many nomadic species are in decline (Runge et al., 2014). Nomadism represents an adaptation to environments where resource availability is low and highly variable in space and time. The survival of nomadic populations may depend on their ability to access dynamically changing resources by displaying long-range movement throughout the year (Mueller et al., 2008). Hence, constraining their ability to move far could greatly impact the abundances of nomadic populations. In Mongolia, the construction of the Trans-Mongolian railway in the 1950s was followed by drastic population declines of gazelles to the west of the railway when their habitat was bisected (Lhagvasuren & Milner-Gulland, 1997). Similarly, the winter mortality of spatially confined Przewalski horses (*Equus ferus przewalskii*) was observed to be higher than in more mobile Asiatic wild ass (Kaczensky et al., 2011). Population densities of Thompson’s gazelle (*Eudorcas thomsonii*) in Kenya, another ungulate considered to be nomadic (Fryxell et al., 2004), decreased following the construction of fences, although decreases in migratory wildebeest and impala in the same area were more dramatic (Said et al., 2016). Partially nomadic pronghorn in North America displayed behavioral avoidance close to wind turbines and reduced habitat use near roads, albeit without immediate negative effects (Milligan et al., 2021).

### Alternative explanations for the decrease in mobility

Many ungulate species adjust their mobility patterns in response to environmental conditions, especially forage availability (Bartlam-Brooks et al., 2013; Merkle et al., 2016), and migration distances often decrease with increasing NDVI (Teitelbaum et al., 2015). Nomadic ungulates exhibit high mobility driven by the scattered and irregular occurrence of resources (Dejid et al., 2019; Mueller et al., 2008), and a decrease in mobility could indicate changes in the overall pattern of forage in space and time. Vegetation greenness in the Eastern Steppe has increased during the last decades (Meng et al., 2020), and a recent climate risk assessment of Mongolia reported that both temperatures and precipitation are predicted to increase under most emission scenarios (World Bank Group & Asian Development Bank, 2021), potentially increasing vegetation productivity, and hence forage availability. In this scenario, the need for continual long-distance mobility could be partially alleviated. Although we found no effect of local NDVI on gazelle displacement, we cannot completely rule out that improving forage conditions might have contributed to the observed decrease in mobility.

### Conservation implications for gazelle in the Eastern Steppe

Maintaining large-scale landscape permeability is critical to conserving wide-ranging ungulates such as gazelle (Nandintsetseg et al., 2019). However, impermeable fences along the Trans-Mongolian Railroad and the Mongolian-Chinese border are already impacting gazelle (Ito et al., 2013; Nandintsetseg et al., 2019), and the proposed construction of additional railroads throughout the Eastern Steppe will likely exacerbate this issue (Batsaikhan et al., 2014). Our findings add an additional perspective to this already concerning outlook: landscape permeability in the Eastern Steppe may be further impacted by increasing traffic intensity, creating semi-permeable barriers to movement. Such barriers have been associated with population declines e.g. in migratory mule deer (Sawyer et al., 2017). Currently, we lack the data to assess whether the decline of gazelle mobility has impacted gazelle population sizes in the Eastern Steppe, as assessments of gazelle abundance have been infrequent and vary in methods and range, with the last published survey likely preceding the onset of potential traffic effects (Olson et al., 2005). Traffic volumes in the sparsely populated Eastern Steppes likely are still far from the level observed in other systems. The decrease in gazelle mobility estimated by our models is relatively low compared to the findings of Tucker et al. (2018), who reported that mammal movements decreased by factor 2-3 in areas of high human footprint. Yet, even if no drastic effects are visible to date, the impact of semi-permeable barriers might change drastically once certain permeability thresholds are reached (Sawyer et al., 2013). The number of registered vehicles in the Eastern Steppe is already consistently increasing (National Center for Road Transport, 2022), suggesting that both traffic and traffic-mediated barrier effects may only grow in intensity.

In its vision for long-term development, the Government of Mongolia has declared to create the transportation infrastructure necessary for an export-oriented economy, and to connect all its settlements with roads until 2050 (State Great Hural, 2020). However, this ambitious goal conflicts with the country’s vision to protect biodiversity and maintain ecosystem services. It also comes at the risk of losing the county’s great herds of nomadic ungulates, which not only provide benefits for regional ecotourism and subsistence hunting, but also serve as a symbol for a vast and undisturbed ecosystem whose people themselves are deeply rooted in their nomadic traditions. To conserve Mongolia’s wide-ranging ungulates, it will be crucial for future infrastructure development policies and land use plans to address the issue of decreasing landscape permeability. We suggest that proposed land-use monitoring efforts already part of the Vision 2050 need to be extended to explicitly encompass the proposed and ongoing road construction projects. To implement scientifically sound policies on environmental protection, long-term assessments of gazelle mobility and abundance need to be continued and supplemented with regular surveys on traffic volumes in core gazelle habitat. This will enable conservationists to look past the high interannual and interindividual variability of movement patterns, pre-emptively identify critical thresholds in landscape permeability and react to barrier effects long before mobility has declined to the point that populations are severely affected. Improving our understanding of how gazelle behaviors are affected by traffic will be crucial to develop mitigation measures for declining permeability. Potentially, barrier effects posed by roads could be alleviated by decreasing the risk perception of vehicles, for instance by prohibiting the use of vehicles for hunting purposes.

## Conclusion

Our study raises concerns on how increasing anthropogenic disturbance threatens large-scale ungulate mobility across the world’s largest mostly-intact grassland, and provides new insights on how nomadic ungulates are affected by semi-permeable barriers. The Mongolian Eastern Steppe is widely known as a land without fences, allowing grand accumulations of ungulates to roam freely through a largely undisturbed ecosystem. However, we documented gradual declines in gazelle mobility in the Eastern Steppe, potentially due to increases in vehicle traffic throughout the steppe and related disturbances such as degraded vegetation around dirt roads. While the effects of this declining mobility on gazelle populations are currently unknown, they may foreshadow significant declines in abundance as traffic and road infrastructure continue to expand and landscape connectivity decreases. Further research will be essential to pinpoint critical thresholds in barrier permeability, and to forecast when these thresholds will be reached.

## Acknowledgements

This research was funded by the German Federal Ministry of Education and Research (BMBF), grant number 01LC1820A. J.M.C. and W.F.F. were supported by NSF IIBR 1915347. This work was partially funded by the Center of Advanced Systems Understanding (CASUS) which is financed by Germany’s Federal Ministry of Education and Research (BMBF) and by the Saxon Ministry for Science, Culture and Tourism (SMWK) with tax funds on the basis of the budget approved by the Saxon State Parliament.

