## Supporting information for "Nomadic ungulate movements under threat: Declining mobility of Mongolian gazelles in the Eastern Steppe"

**A1 Calculating displacements from gazelle movement data with different resolution**

Individual tracking times ranged between 105 and 1797 days, with an average tracking time of 566 days. Gazelle collared in different capture events were tracked at differing temporal resolution: The first dataset consisted of GPS tracking data for 23 gazelles over the years 2007-2013. GPS collars obtained one fix every 25-hour with recurring gaps of 10 days due to the battery life. The second dataset included 22 gazelles with 23-hour fix rate and 5 gazelles with hourly fix rate, monitored from 2014 to 2019 (one individual until 2021). The last dataset was obtained from GPS collars with hourly GPS fix and consisted of GPS tracking data for 12 gazelles over the years of 2019-2021. Because of the different temporal resolutions of the gazelle data, joint analysis of all data was possible only at a few temporal intervals without having to rediscretize the data. We opted to calculate 16-day displacement distances as a measure for long-distance mobility. For the hourly and 25-hourly data, intervals covered exactly 16.0 days (384 hours), while for the 23-hourly data, the closest possible interval covered 16.3 days (391 hours). To account for the difference in time, the 16-day displacement distances calculated from hourly and 25-hourly data were multiplied with a correction factor of 1.018.

**A2 Model selection table for driver models**

**Table 1: Candidate models for average and extreme gazelle mobility.**

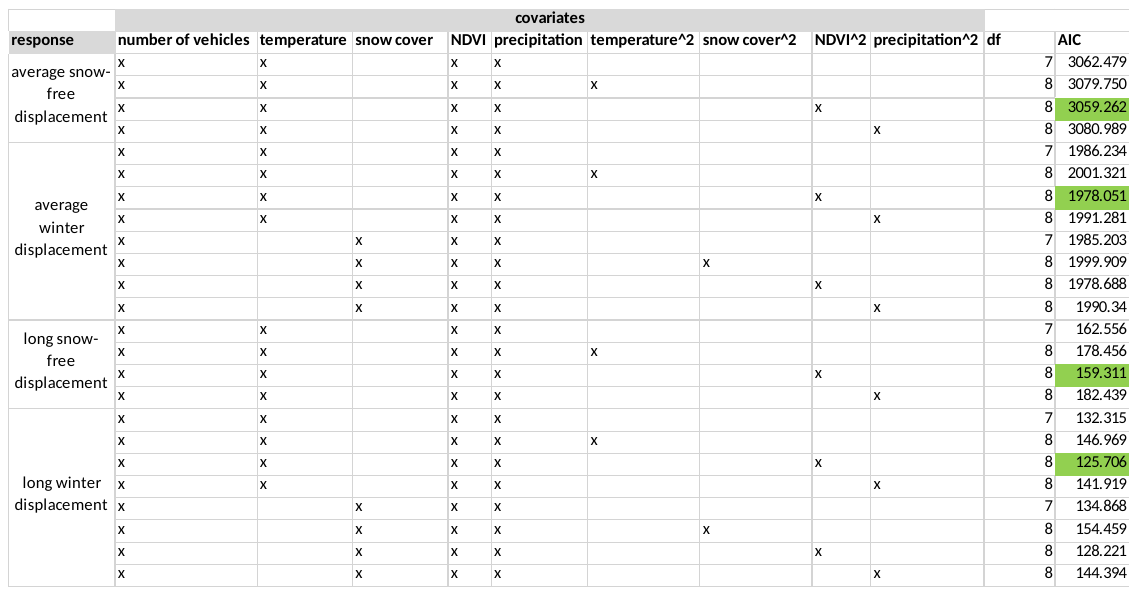

All models included a random intercept for gazelle ID. Including a quadratic effect for NDVI reduced AIC by >2 in each set of models. Including quadratic effects for other environmental predictors never reduced AIC compared to the respective linear-only model. For the winter models, the best model including temperature always outperformed the best model including snow cover. The lowest AIC of each model set is highlighted in green (Table 1).

**A3 Model diagnostics of LMERs**

We assessed model fit using diagnostics of the DHARMa package (Hartig, 2021). Specifically, we used a one-sample Kolmogorov-Smirnov test to assess uniformity of the scaled simulated residuals, and also assessed deviations from homoscedasticity via a test for the location of quantiles. Moreover, we assessed over- and underdispersion using a nonparametric dispersion test via the standard deviation of the fitted vs. simulated residuals and checked for outliers in the simulated residuals based on an exact binomial test with approximate expectations. The diagnostic revealed potentially heteroscedastic errors in three of the models (Table 2, highlighted in red). We subsequently refitted these models as robust linear mixed models via the robustlmm package (Koller, 2016).

**Table 2 Diagnostic tests for LMERs**

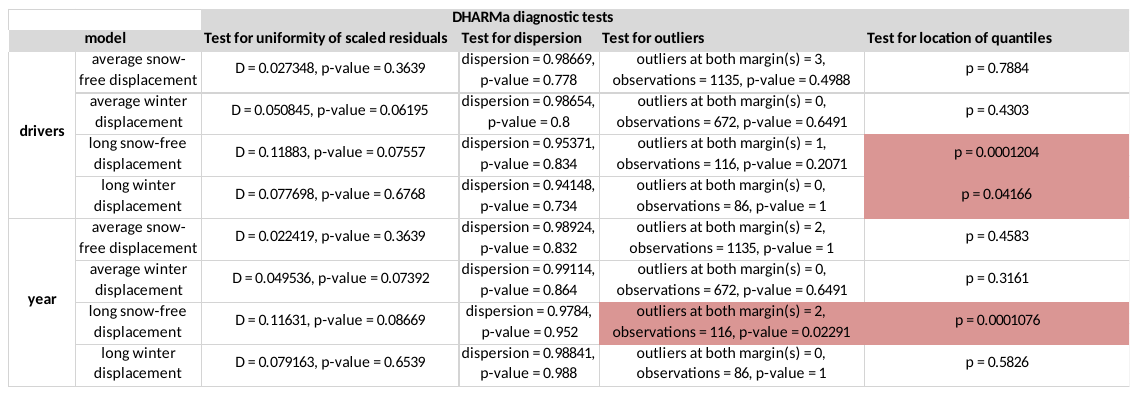

**A4 Spatial eigenvector filtering to reduce spatial autocorrelation in linear models**

We tested for the presence of spatial autocorrelation in the model residuals using the test for Moran’s I in the ape package (Paradis & Schliep, 2019). When we detected significant spatial autocorrelation in the residuals of one of the linear mixed models, we used spatial eigenvectors following Bauman et al. (2018) in order to reduce it. Spatial eigenvectors can be used to control for spatial patterns in regression analysis, aiming to translate the spatial structure of the sampling sites into additional coefficients that can then be entered into a regression model to remove residual spatial autocorrelation (Dray et al., 2006; Bauman et al., 2018). A spatial weighting matrix, consisting of a connectivity matrix of the sampling points and a corresponding weighting scheme, is used to generate eigenvectors by centring rows and columns and standardising (Bauman et al., 2018). If there is significant autocorrelation in the residuals of a regression model, the eigenvectors are entered one by one as additional coefficients, checking for residual autocorrelation after each eigenvector has been incorporated and stopping if Moran’s I in the residuals (MIR) is no longer significant (Griffith & Peres-Neto, 2006). In this MIR approach, the smallest possible subset of eigenvectors is used to preserve as much statistical power as possible for the original predictor variables.

We employed eigenvector selection based on a script that accompanied (Bauman et al., 2018), using the packages spatialreg (Bivand et al., 2021), spdep (Bivand et al., 2022) and adespatial (Dray et al., 2022). Using the starting locations of each displacement step, we built connectivity matrices based on Euclidean distance, identifying the distance up to which significant spatial autocorrelation was present in the residuals of each of the autocorrelated models by plotting correlograms for one-kilometre intervals using the ncf package (Bjornstad, 2022). We identified peaks of significant autocorrelation from the correlograms (basing p-values on 100 permutations) and used the distance-class that showed the first significant peak of autocorrelation as the distance for the connectivity matrix. If this approach failed to reduce spatial autocorrelation to non-significant levels, we incrementally increased the distance until it did. We employed weights that decreased linearly with distance, an approach that was also included in the ones described by Bauman et al. (2018). We then extracted eigenvectors for every autocorrelated model from the corresponding weighting matrix and selected the best subset of them using the MIR approach, following Bauman et al. (2018). After that, we entered those eigenvectors into each corresponding model as additional coefficients, and tested for the presence of spatial autocorrelation in the model residuals as described earlier. This approach successfully reduced residual spatial autocorrelation in all affected models (Table 3), without changing the significance level of the other predictors.

**Table 3 Testing and accounting for spatial autocorrelation**

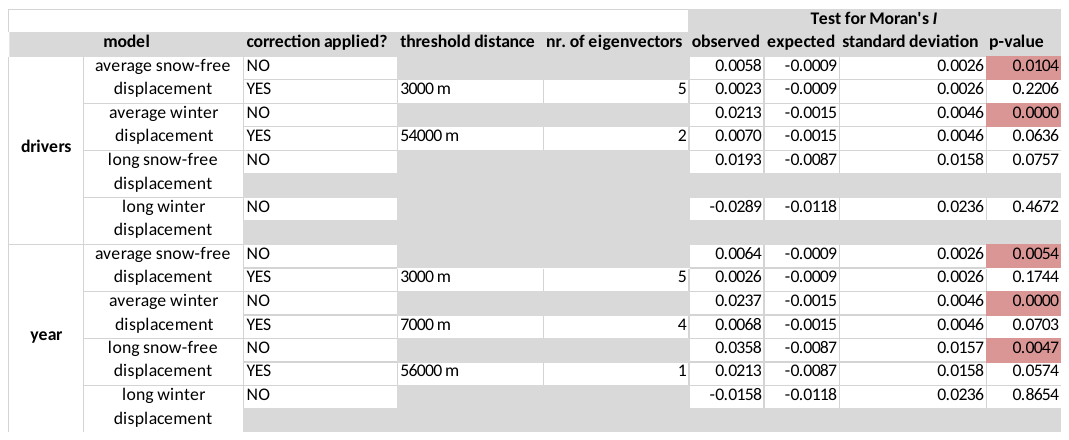

**A5 Model summary tables**

| **Model of average snow-free displacement (drivers)** | | | | | |  |
| --- | --- | --- | --- | --- | --- | --- |
| Fixed effects: |  |  |  |  |  |  |
|  | Estimate | Std. Error | df | t value | Pr(>\|t\|) |  |
| (Intercept) | 3.484 | 0.242 | 593.710 | 14.384 | < 0.001 | *** |
| vehicles | -0.050 | 0.016 | 65.575 | -3.254 | 0.002 | ** |
| temperature | -0.006 | 0.003 | 1123.936 | -1.925 | 0.055 | . |
| NDVI | 2.008 | 1.387 | 1123.999 | 1.448 | 0.148 |  |
| precipitation | 0.002 | 0.002 | 1123.258 | 1.128 | 0.260 |  |
| I(NDVI^2) | -2.819 | 1.921 | 1119.023 | -1.467 | 0.143 |  |
| vec21 | 3.143 | 0.914 | 1102.723 | 3.439 | 0.001 | *** |
| vec40 | 2.512 | 0.900 | 1106.572 | 2.790 | 0.005 | ** |
| vec80 | 2.424 | 0.895 | 1101.732 | 2.707 | 0.007 | ** |
| vec38 | -2.308 | 0.917 | 1123.320 | -2.517 | 0.012 | * |
| vec23 | 2.182 | 0.907 | 1123.858 | 2.406 | 0.016 | * |
| Random effects: |  |  |  |  |  |  |
| Groups | Name | Variance | Std.Dev. |  |  |  |
| ID | (Intercept) | 0.04529 | 0.2128 |  |  |  |
| Residual |  | 0.78461 | 0.8858 |  |  |  |
| Number of obs: 1135 | Groups: ID, 59 | |  |  |  |  |

| **Model of average winter displacement (drivers)** | | | | | |  |
| --- | --- | --- | --- | --- | --- | --- |
| Fixed effects: |  |  |  |  |  |  |
|  | Estimate | Std. Error | df | t value | Pr(>\|t\|) |  |
| (Intercept) | 2.979 | 0.654 | 652.848 | 4.554 | < 0.001 | *** |
| vehicles | -0.017 | 0.020 | 63.359 | -0.856 | 0.395 |  |
| temperature | 0.004 | 0.005 | 645.145 | 0.823 | 0.411 |  |
| NDVI | -4.148 | 7.397 | 662.463 | -0.561 | 0.575 |  |
| precipitation | 0.001 | 0.024 | 663.689 | 0.044 | 0.965 |  |
| I(NDVI^2) | 25.151 | 23.043 | 654.991 | 1.091 | 0.275 |  |
| vec5 | -4.018 | 1.110 | 316.119 | -3.619 | < 0.001 | *** |
| vec7 | 3.325 | 1.119 | 282.830 | 2.970 | 0.003 | ** |
| Random effects: |  |  |  |  |  |  |
| Groups | Name | Variance | Std.Dev. |  |  |  |
| ID | (Intercept) | 0.032 | 0.180 |  |  |  |
| Residual |  | 1.013 | 1.006 |  |  |  |
| Number of obs: 672 | Groups: ID, 59 | |  |  |  |  |

| **Robust model of long-distance snow-free displacement (drivers)** | | | | | |  |
| --- | --- | --- | --- | --- | --- | --- |
| Fixed effects: |  |  |  |  |  |  |
|  | Estimate | Std. Error | df | t value | Pr(>\|t\|) |  |
| (Intercept) | 5.791 | 0.301 | 109.370 | 19.218 | < 0.001 | *** |
| vehicles | -0.078 | 0.018 | 58.418 | -4.461 | < 0.001 | *** |
| temperature | -0.004 | 0.004 | 91.672 | -1.153 | 0.252 |  |
| NDVI | -1.852 | 1.967 | 109.298 | -0.941 | 0.349 |  |
| precipitation | 0.003 | 0.002 | 97.987 | 1.938 | 0.056 | . |
| I(NDVI^2) | 1.659 | 2.693 | 109.951 | 0.616 | 0.539 |  |
| Random effects: |  |  |  |  |  |  |
| Groups | Name | Variance | Std.Dev. |  |  |  |
| ID | (Intercept) | 0.026 | 0.160 |  |  |  |
| Residual |  | 0.105 | 0.324 |  |  |  |
| Number of obs: 116 | Groups: ID, 46 | |  |  |  |  |

| **Robust model of long-distance winter displacement (drivers)** | | | | | |  |
| --- | --- | --- | --- | --- | --- | --- |
| Fixed effects: |  |  |  |  |  |  |
|  | Estimate | Std. Error | df | t value | Pr(>\|t\|) |  |
| (Intercept) | 5.018 | 0.753 | 77.431 | 6.663 | < 0.001 | *** |
| vehicles | -0.049 | 0.021 | 34.359 | -2.358 | 0.024 | * |
| temperature | -0.010 | 0.006 | 79.954 | -1.688 | 0.095 | . |
| NDVI | -1.882 | 9.292 | 78.947 | -0.203 | 0.840 |  |
| precipitation | -0.008 | 0.033 | 79.771 | -0.233 | 0.816 |  |
| I(NDVI^2) | 9.531 | 30.337 | 78.815 | 0.314 | 0.754 |  |
| Random effects: |  |  |  |  |  |  |
| Groups | Name | Variance | Std.Dev. |  |  |  |
| ID | (Intercept) | 0.000 | 0.000 |  |  |  |
| Residual |  | 0.202 | 0.449 |  |  |  |
| Number of obs: 86 | Groups: ID, 42 | |  |  |  |  |

| **Model of average snow-free displacement (year effect)** | | | | | |  |
| --- | --- | --- | --- | --- | --- | --- |
| Fixed effects: |  |  |  |  |  |  |
|  | Estimate | Std. Error | df | t value | Pr(>\|t\|) |  |
| (Intercept) | 60.470 | 18.890 | 61.030 | 3.201 | 0.002 | ** |
| year | -0.028 | 0.009 | 60.930 | -3.025 | 0.004 | ** |
| vec21 | 3.094 | 0.915 | 1107.000 | 3.382 | 0.001 | *** |
| vec40 | 2.586 | 0.897 | 1109.000 | 2.884 | 0.004 | ** |
| vec38 | -2.337 | 0.913 | 1128.000 | -2.560 | 0.011 | * |
| vec80 | 2.438 | 0.895 | 1104.000 | 2.724 | 0.007 | ** |
| vec23 | 2.044 | 0.905 | 1128.000 | 2.258 | 0.024 | * |
| Random effects: |  |  |  |  |  |  |
| Groups | Name | Variance | Std.Dev. |  |  |  |
| ID | (Intercept) | 0.0508 | 0.2254 |  |  |  |
| Residual |  | 0.7832 | 0.885 |  |  |  |
| Number of obs: 1135 | Groups: ID, 59 | |  |  |  |  |

| **Model of average winter displacement (year effect)** | | | | | |  |
| --- | --- | --- | --- | --- | --- | --- |
| Fixed effects: |  |  |  |  |  |  |
|  | Estimate | Std. Error | df | t value | Pr(>\|t\|) |  |
| (Intercept) | 5.691 | 21.025 | 59.518 | 0.271 | 0.788 |  |
| year | -0.001 | 0.010 | 59.384 | -0.132 | 0.895 |  |
| vec25 | -4.899 | 1.050 | 295.904 | -4.666 | < 0.001 | *** |
| vec42 | 3.039 | 1.013 | 661.841 | 3.000 | 0.003 | ** |
| vec102 | -2.764 | 1.017 | 663.795 | -2.717 | 0.007 | ** |
| vec38 | -2.675 | 1.015 | 638.152 | -2.635 | 0.009 | ** |
| Random effects: |  |  |  |  |  |  |
| Groups | Name | Variance | Std.Dev. |  |  |  |
| ID | (Intercept) | 0.016 | 0.125 |  |  |  |
| Residual |  | 1.005 | 1.003 |  |  |  |
| Number of obs: 672 | Groups: ID, 59 | |  |  |  |  |

| **Robust model of long-distance snow-free displacement (year effect)** | | | | | |  |
| --- | --- | --- | --- | --- | --- | --- |
| Fixed effects: |  |  |  |  |  |  |
|  | Estimate | Std. Error | df | t value | Pr(>\|t\|) |  |
| (Intercept) | 106.272 | 18.115 | 52.255 | 5.867 | < 0.001 | *** |
| year | -0.050 | 0.009 | 52.197 | -5.602 | < 0.001 | *** |
| vec2 | -0.821 | 0.365 | 112.671 | -2.248 | 0.027 | *** |
| Random effects: |  |  |  |  |  |  |
| Groups | Name | Variance | Std.Dev. |  |  |  |
| ID | (Intercept) | 0.016 | 0.125 |  |  |  |
| Residual |  | 0.105 | 0.324 |  |  |  |
| Number of obs: 116 | Groups: ID, 46 | | |  |  |  |

| **Model of long-distance winter displacement (year effect)** | | | | | |  |
| --- | --- | --- | --- | --- | --- | --- |
| Fixed effects: |  |  |  |  |  |  |
|  | Estimate | Std. Error | df | t value | Pr(>\|t\|) |  |
| (Intercept) | 74.136 | 24.058 | 36.307 | 3.082 | 0.004 | ** |
| year | -0.034 | 0.012 | 36.291 | -2.889 | 0.006 | ** |
| Random effects: |  |  |  |  |  |  |
| Groups | Name | Variance | Std.Dev. |  |  |  |
| ID | (Intercept) | 0.003 | 0.059 |  |  |  |
| Residual |  | 0.194 | 0.440 |  |  |  |
| Number of obs: 86 | Groups: ID, 42 | |  |  |  |  |

Because robustlmm does not compute p-values, we used the t-statistics of the “robust” rlmer model together with the Satterthwaite degrees of freedom of the “normal” lmer model to determine significance (see e.g. Geniole et al., 2019; Gómez et al., 2022).

We refitted all lmer models without using variable transformations, robust models or autocorrelation corrections to ensure that significant trends did not arise as artifacts of those additions:

| **Uncorrected model of average snow-free displacement (drivers)** | | | | | |  |
| --- | --- | --- | --- | --- | --- | --- |
| Fixed effects: |  |  |  |  |  |  |
|  | Estimate | Std. Error | df | t value | Pr(>\|t\|) |  |
| (Intercept) | 65.722 | 9.515 | 494.558 | 6.907 | < 0.001 | *** |
| vehicles | -2.027 | 0.642 | 52.749 | -3.159 | 0.003 | ** |
| temperature | -0.250 | 0.119 | 1080.444 | -2.102 | 0.036 | * |
| NDVI | 4.466 | 54.047 | 1080.031 | 0.083 | 0.934 |  |
| precipitation | 0.104 | 0.059 | 1080.983 | 1.775 | 0.076 | . |
| I(NDVI^2) | -28.670 | 74.682 | 1079.639 | -0.384 | 0.701 |  |
| Random effects: |  |  |  |  |  |  |
| Groups | Name | Variance | Std.Dev. |  |  |  |
| ID | (Intercept) | 88.57 | 9.411 |  |  |  |
| Residual |  | 1146.18 | 33.855 |  |  |  |
| Number of obs: 1087 | Groups: ID, 59 |  |  |  |  |  |

| **Uncorrected model of average winter displacement (drivers)** | | | | | |  |
| --- | --- | --- | --- | --- | --- | --- |
| Fixed effects: |  |  |  |  |  |  |
|  | Estimate | Std. Error | df | t value | Pr(>\|t\|) |  |
| (Intercept) | 37.570 | 22.126 | 624.747 | 1.698 | 0.090 | . |
| vehicles | -0.775 | 0.652 | 62.150 | -1.189 | 0.239 |  |
| temperature | -0.010 | 0.160 | 615.743 | -0.060 | 0.953 |  |
| NDVI | -119.775 | 251.801 | 631.102 | -0.476 | 0.634 |  |
| precipitation | 0.360 | 0.826 | 630.214 | 0.435 | 0.663 |  |
| I(NDVI^2) | 722.045 | 787.387 | 624.156 | 0.917 | 0.359 |  |
| Random effects: |  |  |  |  |  |  |
| Groups | Name | Variance | Std.Dev. |  |  |  |
| ID | (Intercept) | 40.890 | 6.394 |  |  |  |
| Residual |  | 1159.180 | 34.047 |  |  |  |
| Number of obs: 638 | Groups: ID, 59 |  |  |  |  |  |

| **Uncorrected model of long-distance snow-free displacement (drivers)** | | | | | |  |
| --- | --- | --- | --- | --- | --- | --- |
| Fixed effects: |  |  |  |  |  |  |
|  | Estimate | Std. Error | df | t value | Pr(>\|t\|) |  |
| (Intercept) | 229.431 | 29.242 | 105.708 | 7.846 | < 0.001 | *** |
| vehicles | -6.357 | 1.820 | 52.009 | -3.492 | 0.001 | *** |
| temperature | -0.173 | 0.357 | 87.007 | -0.483 | 0.631 |  |
| NDVI | -337.416 | 191.250 | 105.134 | -1.764 | 0.081 | . |
| precipitation | 0.286 | 0.167 | 92.254 | 1.707 | 0.091 | . |
| I(NDVI^2) | 330.434 | 262.227 | 105.984 | 1.260 | 0.210 |  |
| Random effects: |  |  |  |  |  |  |
| Groups | Name | Variance | Std.Dev. |  |  |  |
| ID | (Intercept) | 383.50 | 19.58 |  |  |  |
| Residual |  | 930.60 | 30.51 |  |  |  |
| Number of obs: 112 | Groups: ID, 46 |  |  |  |  |  |

| **Uncorrected model of long-distance winter displacement (drivers)** | | | | | |  |
| --- | --- | --- | --- | --- | --- | --- |
| Fixed effects: |  |  |  |  |  |  |
|  | Estimate | Std. Error | df | t value | Pr(>\|t\|) |  |
| (Intercept) | 124.631 | 51.308 | 75.499 | 2.429 | 0.018 | * |
| vehicles | -3.776 | 1.463 | 35.532 | -2.581 | 0.014 | * |
| temperature | -0.687 | 0.400 | 77.292 | -1.717 | 0.090 | . |
| NDVI | 14.193 | 631.674 | 77.508 | 0.022 | 0.982 |  |
| precipitation | -0.598 | 2.288 | 77.687 | -0.261 | 0.795 |  |
| I(NDVI^2) | 132.343 | 2065.125 | 77.673 | 0.064 | 0.949 |  |
| Random effects: |  |  |  |  |  |  |
| Groups | Name | Variance | Std.Dev. |  |  |  |
| ID | (Intercept) | 39.520 | 6.286 |  |  |  |
| Residual |  | 937.750 | 30.623 |  |  |  |
| Number of obs: 84 | Groups: ID, 42 |  |  |  |  |  |

| **Uncorrected model of average snow-free displacement (year effect)** | | | | | |  |
| --- | --- | --- | --- | --- | --- | --- |
| Fixed effects: |  |  |  |  |  |  |
|  | Estimate | Std. Error | df | t value | Pr(>\|t\|) |  |
| (Intercept) | 2548.895 | 791.469 | 51.291 | 3.220 | 0.002 | ** |
| year | -1.242 | 0.393 | 51.222 | -3.163 | 0.003 | ** |
| Random effects: |  |  |  |  |  |  |
| Groups | Name | Variance | Std.Dev. |  |  |  |
| ID | (Intercept) | 102.2 | 10.11 |  |  |  |
| Residual |  | 1142.5 | 33.8 |  |  |  |
| Number of obs: 1087 | Groups: ID, 59 |  |  |  |  |  |

| **Uncorrected model of average winter displacement (year effect)** | | | | | |
| --- | --- | --- | --- | --- | --- |
| Fixed effects: |  |  |  |  |  |
|  | Estimate | Std. Error | df | t value | Pr(>\|t\|) |
| (Intercept) | 753.499 | 753.709 | 63.460 | 1.000 | 0.321 |
| year | -0.357 | 0.374 | 63.348 | -0.953 | 0.344 |
| Random effects: |  |  |  |  |  |
| Groups | Name | Variance | Std.Dev. |  |  |
| ID | (Intercept) | 31.240 | 5.590 |  |  |
| Residual |  | 1169.770 | 34.200 |  |  |
| Number of obs: 638 | Groups: ID, 59 |  |  |  |  |

| **Uncorrected model of long-distance snow-free displacement (year effect)** | | | | | |  |
| --- | --- | --- | --- | --- | --- | --- |
| Fixed effects: |  |  |  |  |  |  |
|  | Estimate | Std. Error | df | t value | Pr(>\|t\|) |  |
| (Intercept) | 9472.879 | 2022.093 | 47.953 | 4.685 | < 0.001 | *** |
| year | -4.642 | 1.003 | 47.903 | -4.626 | < 0.001 | *** |
| Random effects: |  |  |  |  |  |  |
| Groups | Name | Variance | Std.Dev. |  |  |  |
| ID | (Intercept) | 377.200 | 19.420 |  |  |  |
| Residual |  | 981.300 | 31.330 |  |  |  |
| Number of obs: 112 | Groups: ID, 46 | | |  |  |  |

| **Uncorrected model of long-distance winter displacement (year effect)** | | | | | | | | | | |  |
| --- | --- | --- | --- | --- | --- | --- | --- | --- | --- | --- | --- |
| Fixed effects: |  | |  | |  | |  | |  | |  |
|  | Estimate | | Std. Error | | df | | t value | | Pr(>\|t\|) | |  |
| (Intercept) | 4974.969 | | 1749.530 | | 37.202 | | 2.844 | | 0.007 | | ** |
| year | -2.418 | | 0.868 | | 37.194 | | -2.785 | | 0.008 | | ** |
| Random effects: |  | |  | |  | |  | |  | |  |
| Groups | Name | | Variance | | Std.Dev. | |  | |  | |  |
| ID | (Intercept) | | 49.160 | | 7.011 | |  | |  | |  |
| Residual |  | | 933.160 | | 30.548 | |  | |  | |  |
| Number of obs: 84 | Groups: ID, 42 | |  | |  | |  | |  | |  |
| **Generalized additive model of the effect of proximity to roads on mobility** | | | | | | | | | | | |
| Parametric coefficients: | | | | | | | | | | | |
|  | | Estimate | | Std. Error | | z value | | Pr(>\|z\|) | |  | |
| (Intercept) | | 3.2202 | | 0.03918 | | 82.18 | | < 0.001 | | *** | |
| season_winter | | -0.36757 | | 0.05217 | | -7.045 | | < 0.001 | | *** | |
| (Intercept).1 | | -0.15972 | | 0.02336 | | -6.837 | | < 0.001 | | *** | |
| season_winter.1 | | 0.06778 | | 0.0396 | | 1.712 | | 0.087 | | . | |
| Approximate significance of smooth terms: | | | | | | | | | | | |
|  | | edf | | Ref.df | | Chi.sq | | p-value | |  | |
| s(log10_distance_to_road) | | 3.676 | | 4.662 | | 73.291 | | < 0.001 | | *** | |
| s(ID) | | 6.89 | | 82 | | 11.441 | | 0.003 | | ** | |
| s(year) | | 4.87 | | 12 | | 12.592 | | 0.006 | | ** | |
| s(x,y) | | 13.254 | | 17.789 | | 67.417 | | < 0.001 | | *** | |
| s.1(log10_distance_to_road) | | 1.763 | | 2.23 | | 2.541 | | 0.273 | |  | |
| Deviance explained = 15.8 | |  | |  | |  | |  | |  | |
| REML = 1490.5 | |  | |  | |  | |  | |  | |
| Scale est. = 1 | |  | |  | |  | |  | |  | |
| n = 1550 | |  | |  | |  | |  | |  | |

| **Additive quantile mixed model on the seasonal differences of distance to roads** | | | | | | |
| --- | --- | --- | --- | --- | --- | --- |
| Quantile |  | Estimate | Std. Error | t value | Pr(>\|t\|) |  |
| 0.9 | (Intercept) | 5931.040 | 263.990 | 22.467 | < 0.001 | *** |
|  | season_winter | -536.480 | 654.900 | -0.819 | 0.415 |  |
|  | s(year) | -558.460 | 296.430 | -1.884 | 0.063 | . |
| 0.75 | (Intercept) | 4692.955 | 129.715 | 36.179 | < 0.001 | *** |
|  | season_winter | -770.704 | 209.984 | -3.670 | < 0.001 | *** |
|  | s(year) | -84.182 | 169.521 | -0.497 | 0.621 |  |
| 0.5 | (Intercept) | 3352.010 | 174.840 | 19.171 | < 0.001 | *** |
|  | season_winter | -512.170 | 207.930 | -2.463 | 0.016 | * |
|  | s(year) | -128.760 | 171.700 | -0.750 | 0.455 |  |
| 0.25 | (Intercept) | 2436.414 | 135.159 | 18.026 | < 0.001 | *** |
|  | season_winter | -438.615 | 148.184 | -2.960 | 0.004 | ** |
|  | s(year) | 12.193 | 161.618 | 0.075 | 0.940 |  |
| 0.1 | (Intercept) | 1805.400 | 85.874 | 21.024 | < 0.001 | *** |
|  | season_winter | -261.439 | 101.161 | -2.584 | 0.011 | * |
|  | s(year) | 123.471 | 89.902 | 1.373 | 0.173 |  |
|  | Observations: 380 | Groups:114 | |  |  |  |

**A6 Model diagnostics of GAMs**

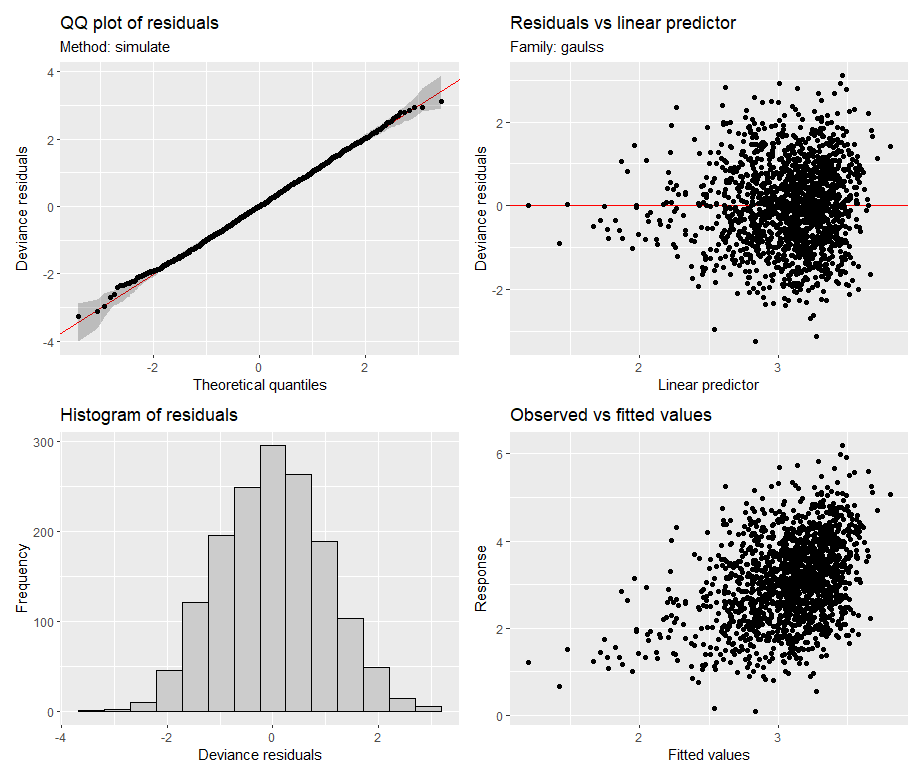

**Figure 1: Diagnostic plots for the generalized additive model of the effect of distance to roads on gazelle displacement.** The employed transformation of the response variable resulted in normally distributed residuals. Residuals appear to be heteroscedastic, therefore we used a Gaussian location-scale model to achieve unbiased p-values in the presense of heteroscedasticity.

Including an interaction between season and distance to road did not decrease Akaike’s Information Criterion (AIC: 4105.993, df: 49.46437) compared to the model without interaction (AIC: 4104.365, df: 44.71698), the interaction was therefore discarded for the sake of parsimony.

After incorporating a spatial smooth of the geographic coordinates, spatial autocorrelation was no longer significant (Test for Moran’s *I*: observed = 0.0034; expected = -0.0006; sd = 0.0022, p-value = 0.0642).
